# Peanut Shell Carbon Nanocellulose Preparation, Analysis and Plausible Role in Regenerative Medicine Using Mesenchymal Stem Cell Line C3H10t1/2

**DOI:** 10.1101/2023.04.19.537474

**Authors:** Kannupriya, Yeshika Bhatia, Seemha Rai, Nidhi Mahajan

**Affiliations:** Center for Stem Cell Tissue Engineering and Biomedical Excellence, Panjab University Chandigarh; Department of Hepatology PGIMER, Chandigarh

## Abstract

Peanut Shell is a major agro industry waste. Cellulose, the most abundant crystalline component of naturally porous peanut shell biomass. Nanomaterial science has actively used peanut shell as a source of nanocellulose. However, there are few reports on the biocompatibility of cellulose nanocrystal based nanocomposites from agro on mesenchymal cell lines. To evaluate how mesenchymal stem cells behave onto this scaffold in vitro, cellulose nanocrystals (CNCs) made from peanut shells were combined with polyvinyl alcohol (PVA) to create tissue engineering scaffolds. SEM images illustrated that for PVA/CNC nanocomposites and clean PVA respectively, increasing the CNC concentration was accompanied with pore size enlargement. The nanocomposite’s three-dimensional porous structure conveyed uneven and intertwined pore structures in additionto the pore distribution. The highest relative crystallinity was observed at 10 wt% of CNCs, according to X-ray diffraction, which also showed that the relative crystallinity of the PVA with 0 and 5 wt% of CNCs was lower than that of neat PVA. In order to confirm the modifications in chemical functional groups, Fourier transform infrared spectroscopy was utilized. The non-cytotoxicity of PVA/CNC_10% was measured for cell viability during an in vitro cytotoxicity test. Additionally, the acquired PVA/biocompatibility CNC’s with the Murine Mesenchymal cell line (C 3H 10T 1/2) demonstrated good cell spreading and adherence to the material surfaces. These results suggest that future research into the in vitro integration of mesenchymal cells with a PVA/CNC scaffold can prove to be a promising candidate for regenerative purposes.

## 1. Introduction

Regenerative medicine is a broad field that includes research on self-healing and tissue engineering as an emerging field of biomedical sciences. Tissue engineering is a discipline of biomedical engineering that seeks to repair, replace and regenerate tissues or organs by using the fundamentals of physics, chemistry and biology to develop effective devices and clinical strategies.

The process of tissue engineering involves cells and biomolecules combined with scaffolds [1]. Scaffolds are 3D constructs which can be synthetic or natural and deliver the physicochemical and mechanical maintenance for *in vitro* ECM formation, which further gets degraded, resorbed and metabolized when implanted *in vivo* [2]. Scaffolds are typically made up of polymeric biomaterials which provide structural support for cell attachment and the development of tissue.There are a variety of choices when selecting a scaffold for tissue engineering. Instinctively, the best scaffold for engineered tissue should mimic the ECM of the target tissue, associated with its biocompatibility, biodegradability, its architecture and mechanical properties [3]. For the production of scaffold based on biological material such as collagen and a variety of proteoglycans such as cellulose, chitosan and alginate-based substrate. Cellulose is the most abundant and well known natural polymer available and has been used to produce diverse products ranging from industrial to medical applications such as fibers, haemodialysis and blotting membranes [4,5]. Cellulose scaffolds are one of the most suitable materials for cell proliferation and differentiation because of their mechanical and physical properties [6]. A possible source of several useful chemicals is groundnut shell, a byproduct of the significant leguminous crop groundnut. Groundnut shells account for approximately 20% of dried groundnut pod by weight and are the major waste of groundnut industry [7,8]. Majorly used for composting kitchen waste, animal feed, utilized for burning for producing energy and now days they are being converted into various bio-products such as biodiesel, bioethanol, hydrogen production etc. [7,8,9] Due to the readily available natural antioxidants in groundnut shells, which contain antioxidant, antibacterial and inhibitory properties, many chemists and nutritionists have become interested in them [10,11]. Moreover, due to its high cellulosic and hemicellulosic content, it can be use dfor scaffolding in tissue engineering [12,18]. As it is used to treat coughs, lower blood pressure, and clear mucus from the lungs, it is receiving more attention in the therapeutic and pharmaceutical fields [14,15]. Consumer demand for groundnut shell products is rising in the health and cosmetic industries as a result of safety and health-promoting concerns [16].

## 2. Materials and methods

The groundnut shells were collected after the process of blanching the groundnuts collected from a local vendor. Pure grade methanol was used for the soxhlet process for extraction of groundnut shells. For cell culture, the mesenchymal C_3_H_10_t_1/2_ cell line was used. For culturing, Dulbecco’s Modified Eagle’s Medium (DMEM), Fetal Bovine Serum (FBS), Trypsin, and Ethylene Diammine Tetracaetato (EDTA) were used for culturing. Cell culturing of C_3_H_10_t_1/2_ cells was performed in full growth media for further assays such as Trypan blue and MTT.

### 2.1. Isolation of cellulose

The groundnut shells were washed thoroughly with distilled water and then sundried. The sundried groundnut shells were grounded into a fine powder and sieved through a 150mm mesh. 25g of sieved groundnut shell powder was treated with sodium hydroxide (0.5M) for two hours at 90 C with continuous stirring. The obtained dark slurry was filtered and washed with distilled water several times. The washed residue was refluxed with a solution containing 20% (v/v) of Nitric Acid (HNO_3_) in ethanol three times, and the color changed from brown to yellow. The mixture was then filtered and washed with distilled water to neutrality. The yellow-colored residue was then bleached with Sodium Hypochlorite (NaOCl)to an off-white color. The cellulose obtained was lyophilized for 24 hours to get a fine powder of cellulose [17,18].

### 2.2. Preparation of nanocellulose crystals (CNC)

5g of cellulose powder was hydrolyzed for 75 minutes in 60% (w/v) sulfuric acid at 45 degrees Celsius with continuous stirring at 500rpm on a magnetic stirrer. After that, cold distilled water was added to quench the reaction, and the slurry was washed with distilled water using repeated centrifugation (10000–12000rpm for 15 min) and a pH of 7 was maintained. After that, sonication was conducted in an ice bath for 10 minutes. The obtained aqueous suspension was stored in are frigerator at 4°C for further use. [17,18,19,20].

### 2.3. Preparation of bioscaffold

The 5wt% PVA solution was prepared in distilled water at 70°C for 10 hours with continuous slow stirring. Further 0, 5, and 10% on CNCs were added and stirred for more than 3 hours to thoroughly mix the components. Then the suspension was sonicated for 10 minutes to remove bubbles. The bioscaffold were cast on a 24-well plate and frozen at -80°C for 12 hours and freeze-dried under vacuum for 3 days. [20,29]

### 2.4. Characterization methods

#### 1) Morphological analysis

##### Transmission Electron Microscopy (TEM)

Transmission electron microscopes (TEM) are types of microscopes that provide an image that is greatly enlarged while viewing specimens using an electron particle beam. TEMs have a 2 million-fold magnification capacity. Consider how little a cell is to gain a better understanding of how tiny that is. It is understandable why TEMs have grown to be so important in the biological and medical disciplines. Here, a 40-120kV operating voltage Hitachi (H-7500) sample was negatively stained.

##### Field-emission scanning electron microscopy (FE-SEM)

Electronic microscopy using field emission scanning Instead of using light, a FESEM microscope uses electrons, which are particles having a negative charge. A field emission source releases these electrons. By moving in a zigzag motion, electrons scan the thing. The surface of whole or fractioned objects can have very minute topographic characteristics that can be visualised using a FE-SEM. This method is used by scientists studying biology, chemistry and physics to examine structures that could be as small as 1 nanometer (= billionth of a millimetre). For instance, organelles and DNA material in cells, synthetic polymers, and coatings on microchips can all be studied with the FE SEM. The surface morphology of PVA/ CNC_0%, PVA/CNC_5%, and PVA/CNC_10% bioscaffold was examined using a field-emission scanning electron microscope (JSM-6100) at a 15V acceleration voltage. For analysis, samples were cut perpendicular to the height of the cylindrical scaffold using a sharp blade and images were taken at 100X, WD 11.

#### 2) Fourier-transform infrared spectroscopy

Due to its exceptional mix of sensitivity, flexibility, specificity and resilience, Fourier transform infrared (FTIR) spectroscopy is a tremendously popular technique today. It is now one of the most frequently used analytical techniques in science and can handle solid, liquid, and gaseous analytes. Although FTIR has some known drawbacks, such as a relative intolerance of water and sensitivity to the physical characteristics of the analysis matrix, it is still widely used in a variety of fields, including the food and beverage industry, engineering, environment, pharmaceuticals, biomass, and clinical settings. FTIR spectroscopy was used to evaluate the structural variations and intermolecular interactions among the components of the bioscaffold. The samples were analysed attenuated total reflectance accessories over the range of 400– 4000cm^− 1^.

#### 3) X-ray diffraction (XRD) analysis

Understanding the three-dimensional structure of biological macromolecules, such as DNA and proteins, is essential to comprehending how life functions. The primary technique of structural biology, the field of biology that investigates the structure and spatial organization of biological macromolecules, is known as biological crystallography. It is based on the investigation of X-ray diffraction by crystals of macromolecules. The specimens were scanned at 2 θ

### 2.5. Cytotoxicity

The cytotoxicity of samples was measured using the MTT assay. The MTT reagent (3-(4,5-dimethylthiazol-2-yl)-2,5diphenyl-2H-tetrazolium bromide) is a mono-tetrazolium salt with thre earomatic rings, two phenyl moieties, and one thiazolyl ring surrounded by a positively charged quaternary tetrazole ring core containing four nitrogen atoms. When MTT is reduced, the central tetrazole ring is broken, creating the violet-blue, water insoluble chemical known as formazan [16].Due to its positive charge and lipophilic composition, the MTT reagent, which is converted to formazan by metabolically active cells, can pass through the mitochondrial inner membrane and cellmembrane of live cells. 5000 cells were seeded on a96-well plate and full growth media was added.Then, the bioscaffold were equally cut into the same dimensions and placed in a 96-well plate where cells are seeded and incubated for 242 hours in a CO_2_ incubator at 372 degrees C. After incubation, MTT reagent was added and absorbance was read at 570nm using a microplate reader. *Relative viability*= *OD of sample* ÷ *OD of control* ×100

### 2.6. Cell count

A population’s percentage of healthy, alive cells is known as its cell viability. Cell viability assays are used to assess the general wellbeing of cells, improve experimental or culture settings, and gauge cell survival after exposure to substances, such as during a drug screen. Cell viability assays frequently evaluate metabolic activity, ATP content, or cell proliferation to give a readout of the health of the cells. Cell toxicity assays, which give a readout on signs of cell death like a loss of membrane integrity, can also be used to measure cell viability. Cell viability and cell toxicity tests are crucial techniques for evaluating cellular reactions to potential experimental substances. Trypsinized cells were pelleted by centrifugation (4000 rpm for 4 minutes at 4 C) and re suspended in 50 uL of growth media. 20uL of cell suspension was mixed with 380uL of count viability volume and loaded onto the Muse Cell analyzer (MCH 100101) and cells were analyzed. *Cell viability=total number* of *viable cells ÷total cells* (*live + dead*) *×* 100

### 2.7. DAPI Staining

The blue-fluorescent DNA stain DAPI (4′, 6-diamidino-2-phenylindole) exhibits a 20fold increase in fluorescence when bound to AT sections of dsDNA. It is frequently used as a nuclear counterstain in fluorescence microscopy, flow cytometry, and chromosomal staining. It is stimulated by the violet (405 nm) laser line. Similarly, here the medium was aspirated from the culture vessel. A sufficient volume of fixative was added to cover the scaffold and stood for 10 minutes at room temperature. Then fixative was removed and 1 ml of working stain solution was added and kept for 30 minutes in dark. Further stain was removed and a section was cut from the scaffold and visualized under a fluorescentmicroscope at 10X [22].

## 3. Results and discussion

With the increasing advances in Tissue Engineering increased the requirement of materials that can be used for tissue reconstruction and scaffolds, cells and biomolecules are basic requirement for tissue reconstruction. In the present study, abiocompatible biodegradable scaffold has been created which is very much cost effective from Groundnut shells. Groundnut shells account for approximately 20% of dried groundnut pod by weight and are the major waste of groundnut industry. The scaffold activity of nanocellulose has been reported in a few studies [19, 20]. However, few studies have examined the proliferation of cells of different cell line [20]. A significant component of tissue engineering is the creation of scaffold for tissue repair, at the forefront of current research, tissue engineering strategies for scaffold formation with cellulose-based materials. In the fields of biomedical engineering, micro-and nanocellulose-based materials are at the cutting edge of scientific advancement. Because of its accessibility and toxicity characteristics, cellulose has garnered a lot of attention in the scaffolding industry. The present study investigates the scaffold activity of nanocellulose from peanut shells by studying the pore size, crystallinity and various modifications in the functional groups. PVA biocomposites with different cellulosic concentrations were made such as 0%, 5%, and 10% respectively. A few studies have reported the proliferation of different cells such as L929 cells [20] on the surface of the cellulose-derived nanocellulose from various sources. In the present study, it was investigated that the cytoxicity of C_3_H_10_t_1/2_ cells on the bioscaffold of peanut shell isolated cellulose. As per FTIR, these peaks in bioscaffold showed varying intensities, indicating changes in the crystallization of PVA as a result of CNC addition, forming hydrogen bonds between PVA and the CNCs. As a result, the FTIR measurements indicated that theCNC and PVA matrix had chemical interactions [19,20,22,23]. In this study we also investigated the adherence and proliferation of C_3_H_10_t_1/2_. Following a 7day PVA/CNC 10% treatment, DAPI staining of the murine mesenchymal stem cells revealed the existence of adherent and proliferating C _3_H _10_ t _1/2_ cells. In our study, we also studied cell viability. To do this, the MTT assay was used to gauge C3H10t1/2’s viability. Utilizing XRD, it was possible to control how the characteristics of a polymeric crystalline structure changed the sameresult was also seen in study carried out by Lam NT(2017) [19,20,22,23]. When making bioscaffold, specific changes to the chemical functional groups were detected using FTIR spectroscopy. The biological response that determines the outcome of tissue restoration heavily depends on the scaffold’s architecture. SEM was used to study the morphology of the cross-section porous bioscaffold.

### 3.1. Process of bioscaffold formation

Fig. 1A and 1B shows steps for the formation of bioscaffold. Fig. 1B(a) is the groundnut shell powder and then it was treated with NaOH solution for 2 hours at 95°C in fig. 1B(b), de-lignified cellulose was obtained in brown color fig. 1B(c) and in fig. 1B(d) acid reflux was done to obtain yellow cellulose in fig. 1B(e) and further bleached to white in fig. 1B(f). The obtained cellulose was acid hydrolyzed to obtain cellulose nanocrystals in fig. 1B(g) and then it was mixed with PVA and casted on 24-well plate fig. 1B(h) and freeze dried to obtain bioscaffold on fig. 1B(i). The cellulose was chemically extracted from washed, sundried Groundnut shell powder by treating with sodium hydroxide then acid refluxed and at-last bleached to produce white cellulose and same process was followed by Kumar A (2014) and Punnadiyil (2016) in sugarcane baggase and Groundnut shells and produced cellulose. The cellulose was then converted into cellulose nanocrystals from the methodology used by Lam NT (2017), Kumar A (2014) and Punnadiyil (2016) in their work done with sugarcane and groundnut.

**Fig. 1.**
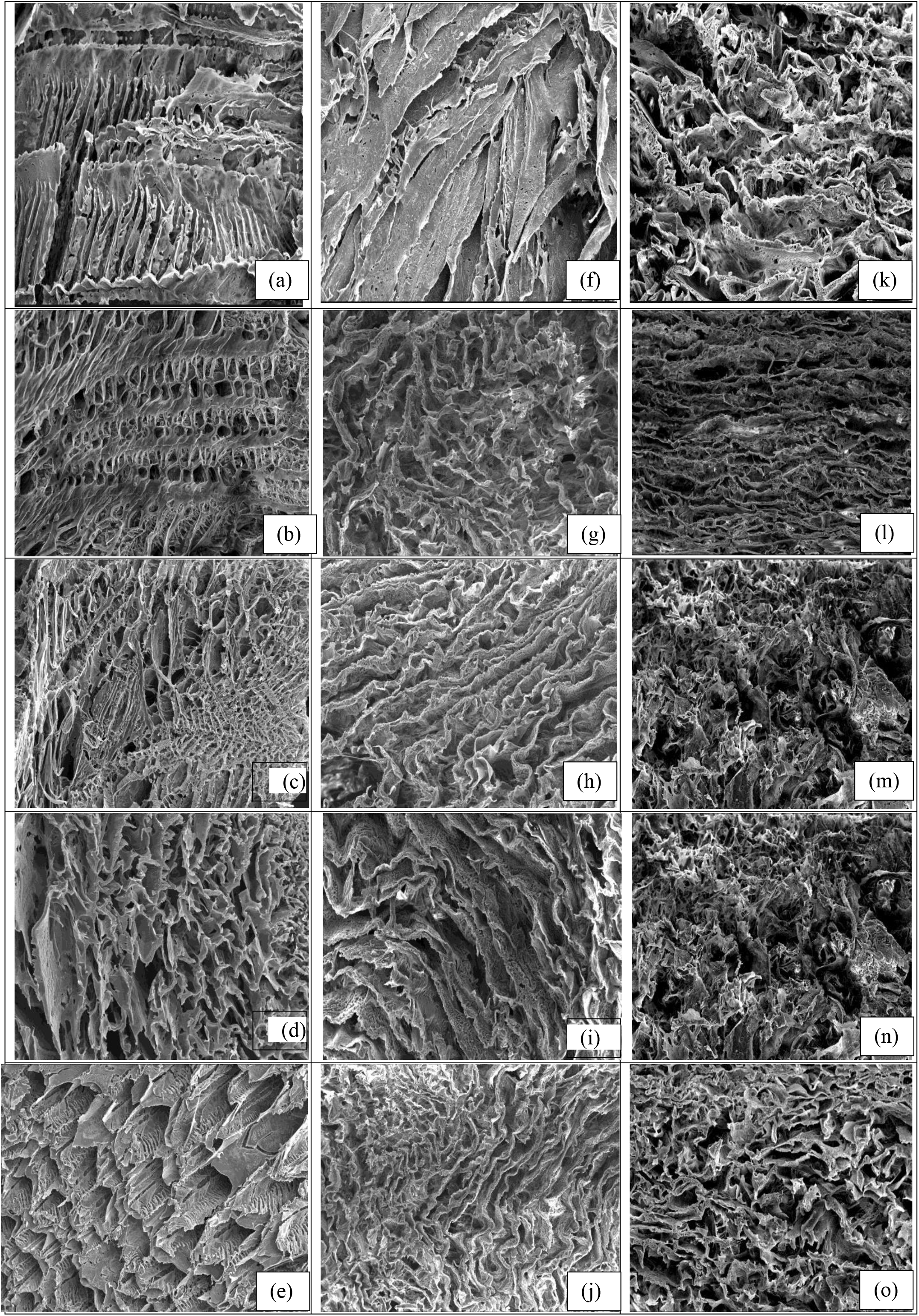
A: Flow chart depicting the bioscaffold formation. Fig. 1. **(A)** Flow chart depecting the bioscaffold formation **(B)** Steps followed for bioscaffold formation (a) Groundnut shell powder, (b) NaOH treatment, (c) Delignified powder, (d) acid reflux, (e) acid refluxed powder, (f) bleached cellulose, (g) CNC, (h) fabricated and (i)freeze dried scaffold.

### 3.2. Morphology study

#### 1) Cellulose nanocrystal analysis

A beam of electrons is transmitted through a specimen in Transmission Electron Microscope (TEM) to produce a picture. The interaction between the beam’s electrons and the sample results in a variety of signals that can be used to learn more about the surface’s topography and and composition. Cellulose nanocrystals were obtained by acid hydrolysis and were obtained as a white colloidal suspension. Fig. 2 shows a SEM micrograph of CNCs with a pine-like shape. [19,22] The same results were seen when cellulose nanocrystals was obtained from sugarcane baggase by Lam NT (2017). Cellulose nanocrystals were obtained by acid hydrolysis and were obtained as a white colloidal suspension. Fig. 2 shows a SEM micrograph of CNCs with a pine-like shape. [19,22] The same results were seen when cellulose nanocrystals was obtained from sugarcane baggase by Lam NT (2017).

**Fig. 2.**
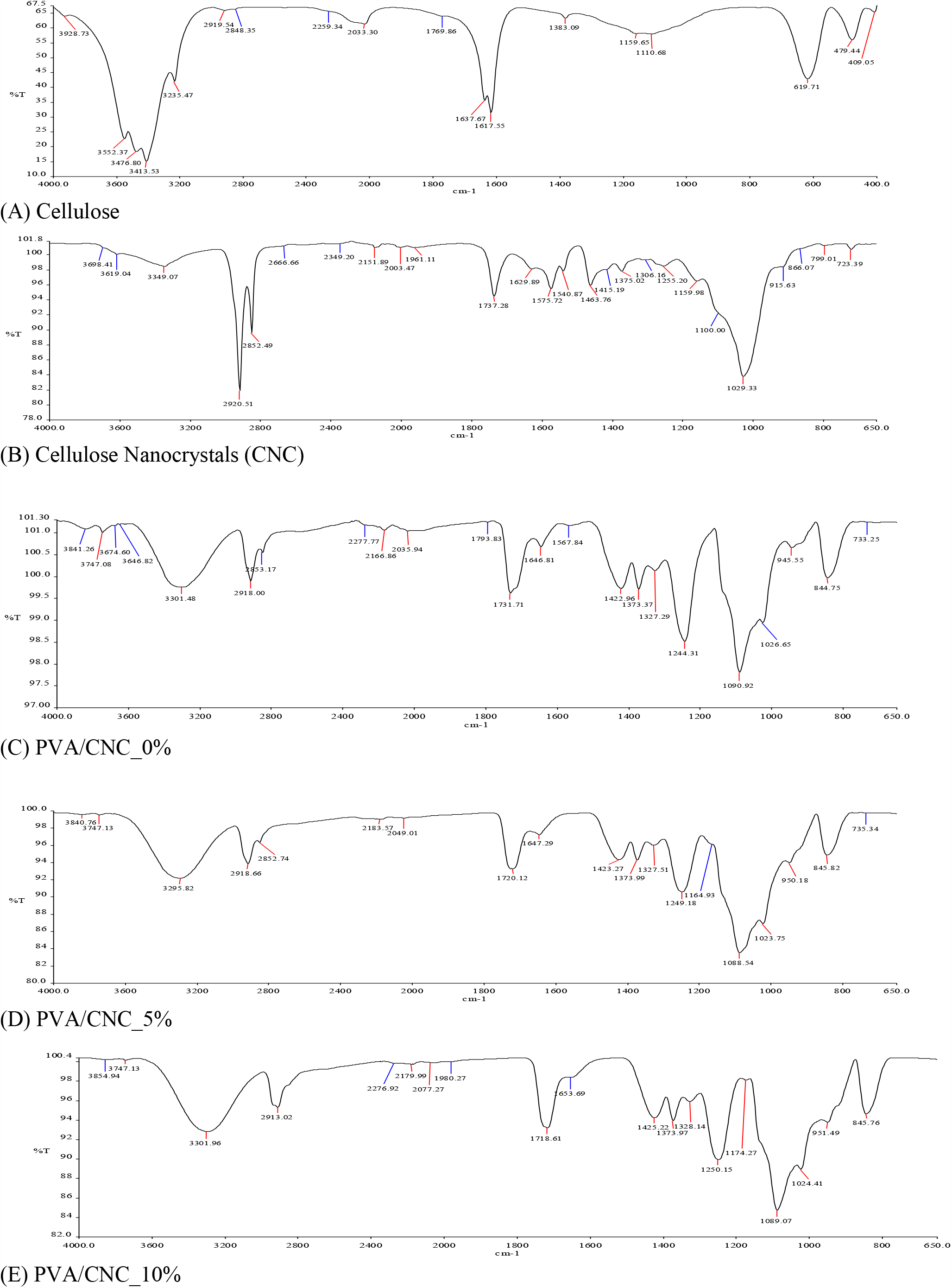
TEM images of cellulose nanocrystals. (CNC) (a) and (b) are TEM images of cellulose nanocrystals formed after acid hydrolysis of cellulose. (a) TEM images of CNC at 200nm and (b) TEM images of CNC at 500nm.

#### 2) FE-SEM analysis

The architecture of a scaffold is a key part of the biological response that determines the success of tissue reconstruction. The morphology of the cross-section porous bioscaffold was observed by SEM as shown in fig. 3. It was observed that the porous structure was affected by the addition of cellulose bioscaffold. On the other hand, the cross linkages among the hydroxyl groups of cellulose bioscaffold were seen to increase by increasing the cellulose nanocrystals content. SEM imaging of obtained PVA/CNC bioscaffold was done which showed that with adding CNC content the pore size is decreased and more circular voids are seen in scaffold, the same morphology was obtained with SEM images of bioscaffold obtained by sugarcane baggase Lam NT (2017).

**Fig. 3.**
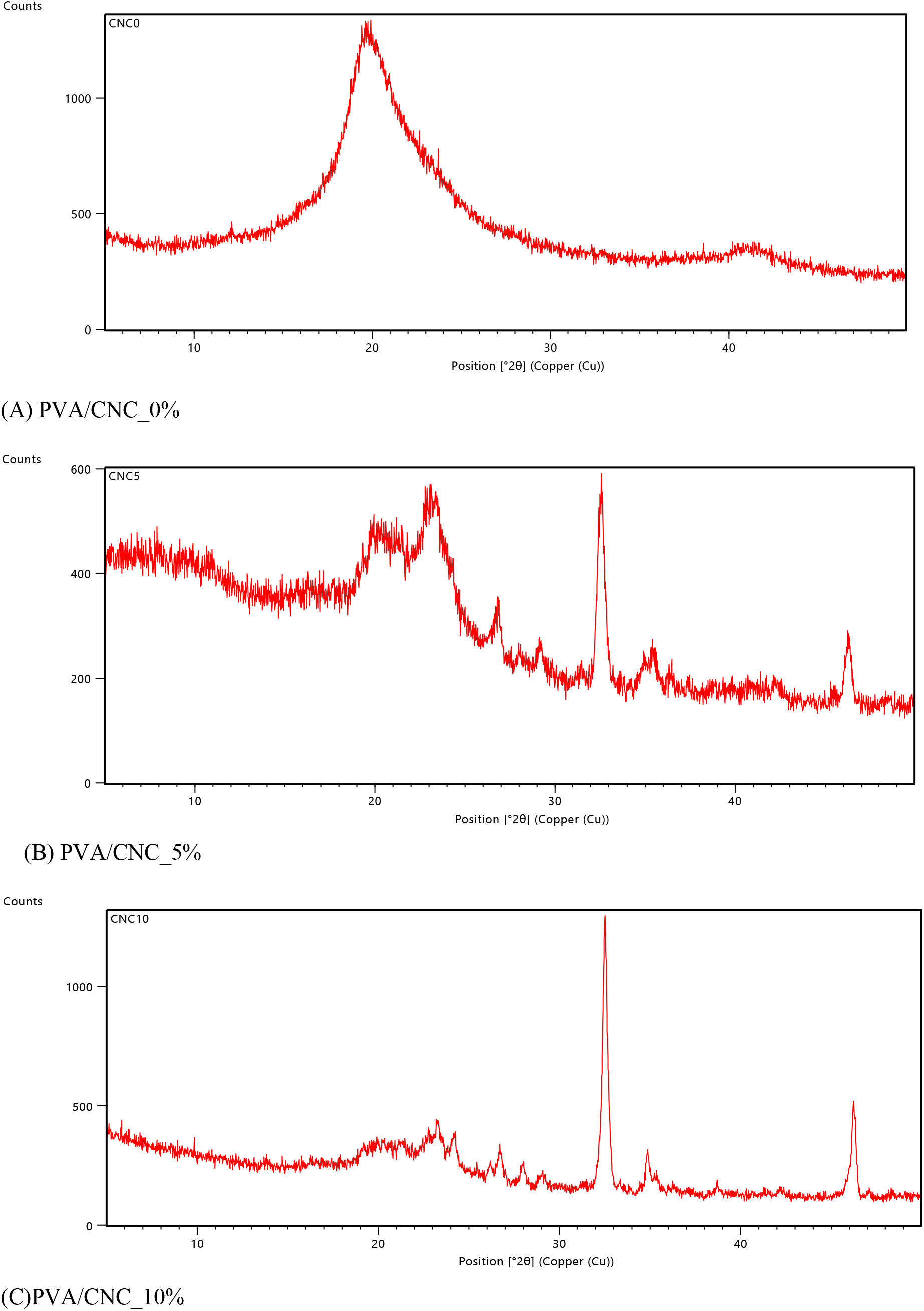
FE-SEM micrographs of (a)-(e) PVA/CNC_0%; (f)-(j) PVA/CNC_5%; (k)-(o)PVA/CNC_10%

##### 3.3. FTIR spectroscopy

FTIR spectroscopy was used to identify certain alterations in the chemical functional groups during the fabrication ofbioscaffold. FTIR gives the resultant graphical representation, which exhibits the typical shape of elasto-plastic foam. FTIR spectra of cellulose, cellulose nanocrystals (CNC), PVA/CNC_0%, PVA/CNC_5%, and PVA/CNC_10% nano-composite scaffolds in the 4000 to 400cm-^1^ spectral range are shown in Fig.s 4(a) to 4(e). FTIR gives the resultant graphical representation, which exhibits the typical shape of the elastoplastic foam. The FTIR absorbance curves are the mean values of the compression modulus. In fig. 4(a), the absorption band of cellulose can be seen at 3200– 4600cm^-1^, which is evidence of the presence of a large number of many types of hydrogen bonds due to –OH groups. Fig. 4(b) shows the absorption band in the 2200–3300cm^-1^ region can be seen in cellulose nanocrystals (CNC), which depicts O-H bending of adsorbed water. And absorption in the region 970-1250 cm^1^ shows the presence of C-O bonds. In FTIR spectra of bioscaffold, fig.s 4(c) to 4(e), the broad absorption band in the 3200– 3550cm^-1^ region indicates strongly absorbed and bound water in bioscaffold, depicting the OH bond. Absorption in the range of 2850–3000 cm^-1^ shows the formation of an alkane bond. An absorption peak of saturated aldehyde can be seen in the region of 1720–1740 cm^-1^. Skeletal C-C vibrations can be seen in regions 1300–700 cm-1 [19,20,22,23] The changes in functional group of cellulose, cellulose nanocrystals, PVA/CNC composites can be seen by FTIR and being rich many different hydrogen bonds cellulose have shown broad peak in that region, cellulose nanocrystals have shown O-H bending of absorbed water during acid hydrolysis. PVA/CNC has shown absorbed and bound water in nanocomposite alkane bond and the similar data was obtained in FTIR done by Punnadiyil (2016) and Lam NT (2017). The decreased/increased intensity of these peaks in bioscaffold suggested changes in in the crystallization of PVA due to CNC addition creating hydrogen linkages between PVA and the CNCs. Consequently, the FTIR results suggested the occurrence of chemical interactions between CNC and PVA matrix.

**Fig. 4.**
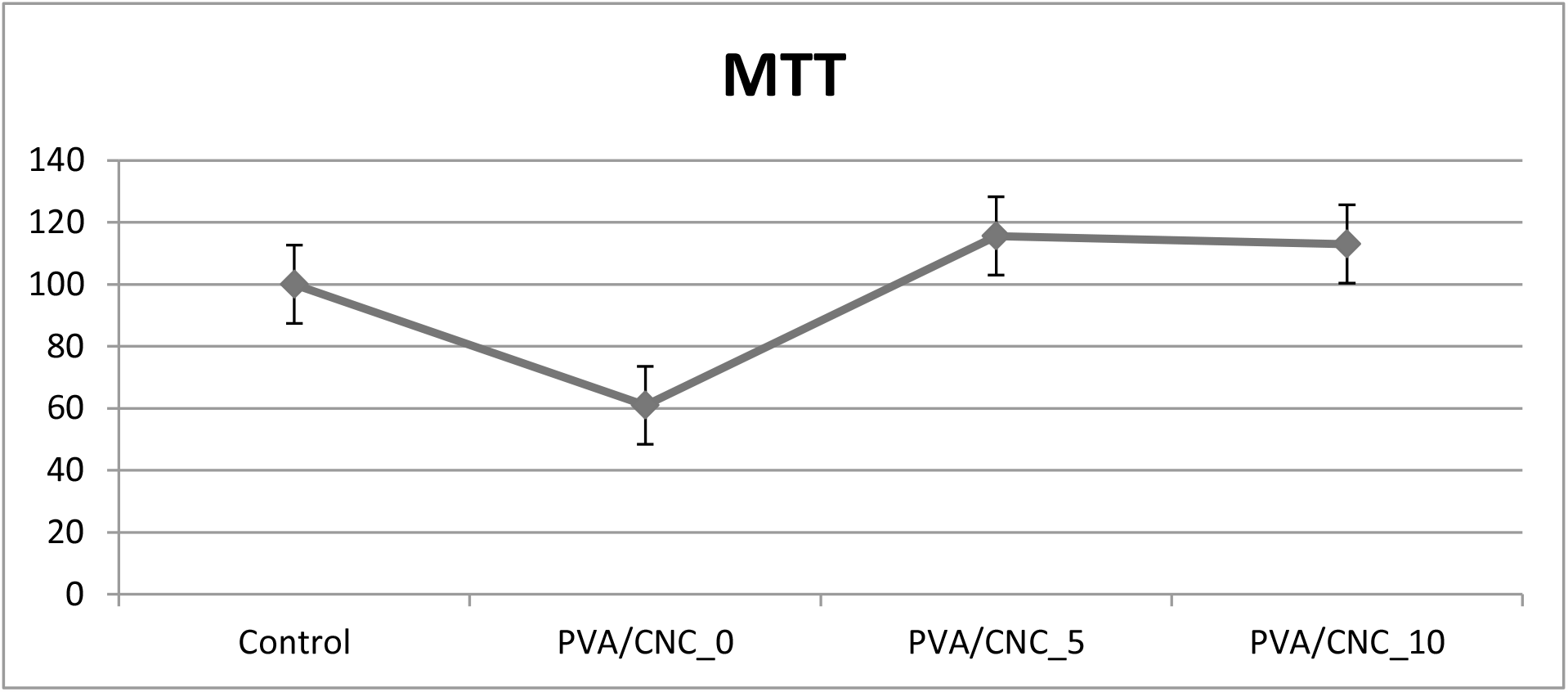
FTIR spectra of (A) Cellulose; (B) Cellulose Nanocrystals (CNC); (C) PVA/CNC_0%; (D) PVA/CNC_5%; (E) PVA/CNC_10%

##### 3.4. XRD Pattern

XRD was exploited to direct the changes in the features of a polymeric crystalline structure. The graphs employed by XRD that is, intensity versus theta, are basically used to find the nature of material. The sharp peak indicates that the material is crystalline; similarly, the broader peak indicates that the material is polycrystalline; and no peak indicates that the material is amorphous. The crystallinity percentage of the bioscaffold is presented in fig. 5. The spectral patterns analyzed by XRD suggested a semi-crystalline nature of the fabricated PVA/CNC scaffolds [19,20,22,23]. The crystallinity of different PVA/CNC compound was measured by X-ray diffraction (XRD) and the results have shown that PVA/CNC_0% had no sharp peak depicting amphoteric nature of composite, PVA/CNC_5%, PVA/CNC_10% have shown sharp peaks showing semi-crystalline nature of bioscaffold and PVA/CNC_10% have shown more peaks is due to more amount of CNC.

**Fig. 5.**
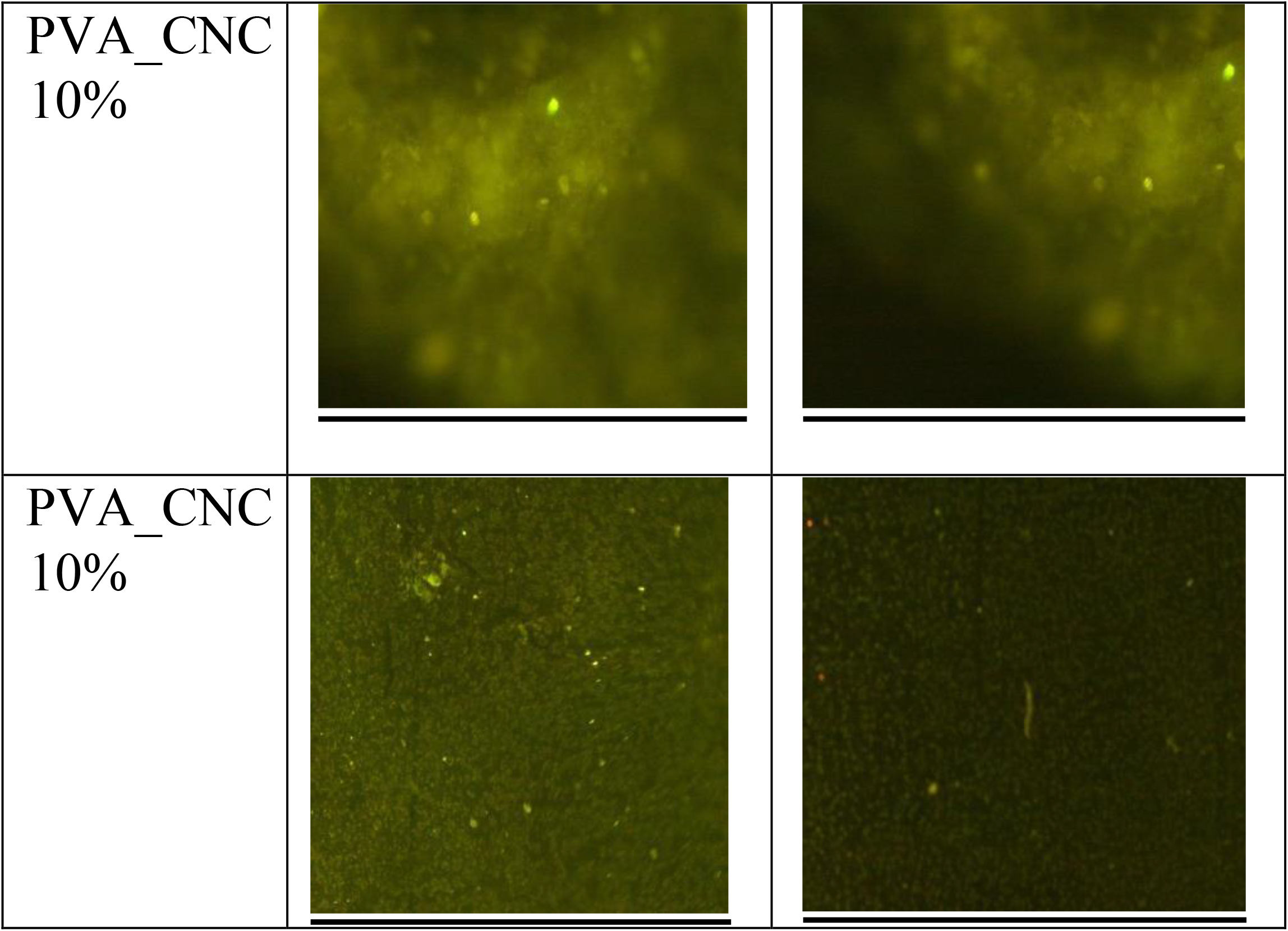
XRD patterns of (A) PVA/CNC_0%; (B) PVA/CNC_0%; (C) PVA/CNC_10% scaffolds. All the scaffolds exhibit semi-crystalline nature.

**Fig. 6:**
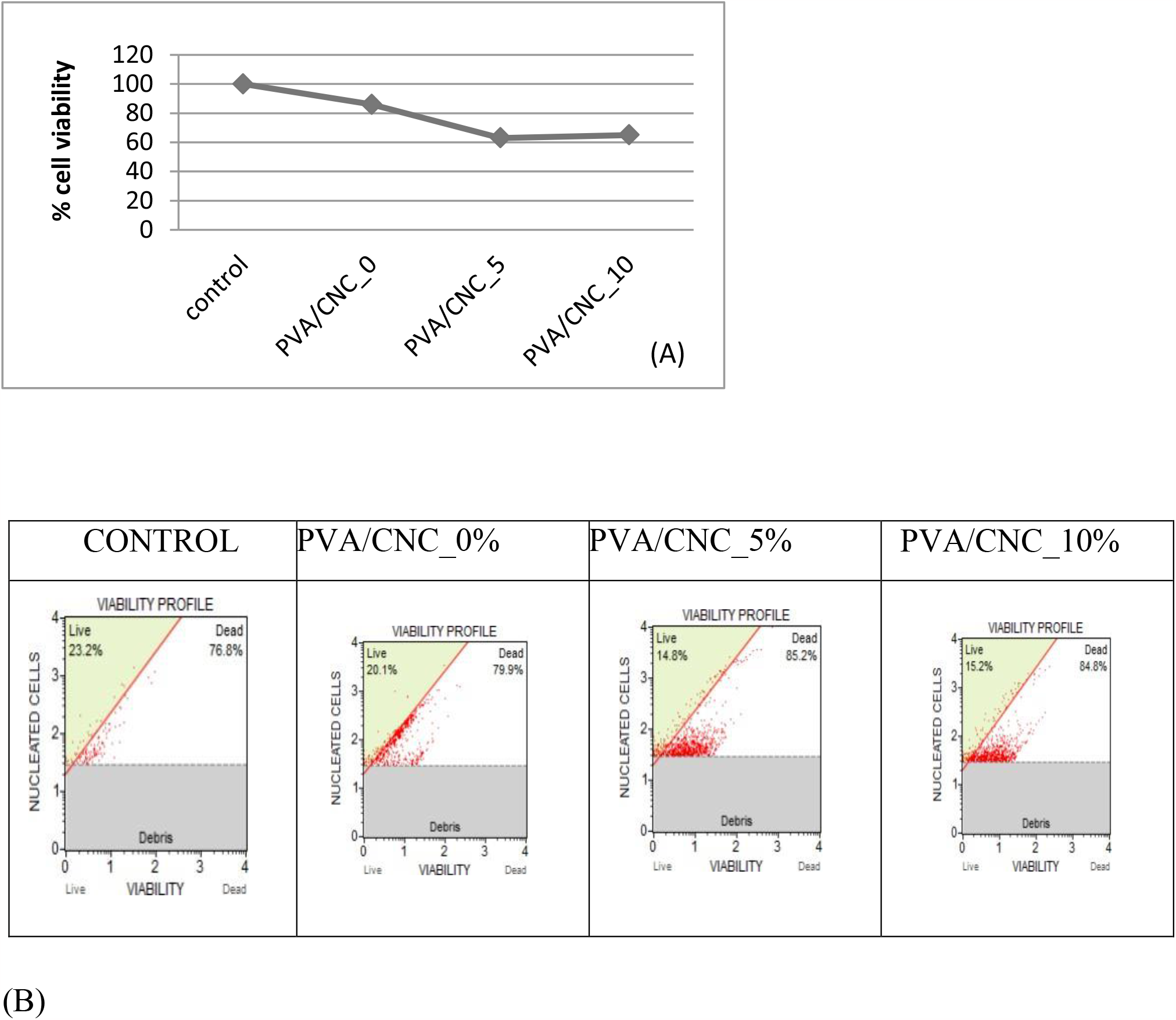
Results of MTT performed. control is taken to be 100% and relative percentage viability was measured. Treatments with PVA/CNC bioscaffold have shown to increase cell viability.

##### 3.5. Cell culture of mesenchymal cells C3H10t1/2

The C_3_H_10_t_1/2_ cell line was developed in 1973 from C_3_H mouse embryos that were 14 to 17 days old. These cells resemble mesenchymal stem cells interms of functionality and exhibit a fibroblastic shape in cell culture. This cell line serves many purposes and types in tissue engineering. The cells were cultured in nutrient media: Dulbecco’s modified Eagle’s medium supplemented with foetal bovine serum and antibiotics. The medium alone was used as a control. The culture medium was changed every 3 days. The relative crystallinity of bioscaffold was calculated using Origin 8.0 software using the surface method [23].

##### 3.6. Cell viability

Before the *in vitro* testing of any implantable biomaterial (bioscaffold), it is crucial to evaluate the biocompatibility of the biomaterial in terms of the non-toxic concentration in the cell line. For this, the MTT assay was employed to estimate the viability of C _3_H _10_ t _1/2_. After the cells were incubated for 24h, there were statistically insignificant or significant (choose between two) differences in cell morphology. The viability threshold for cytotoxicity analysis was calculated by the MTT assay where cell cytotoxicity in case PVA/CNC_10 was found to be maximum as shown in fig. 4. The viability of the control was set at 100% [21]. To check biocompatibility Mesenchymal stem cells (C_3_H_10_t_1/2_) were grown on bioscaffold and cell viability was measured by MTT after 24 hours of cell plating on bioscaffold. Treatment with PVA/CNC bioscaffold have shown to increase cell viability and enhanced cell growth. In work done by Alhosseini (2015) and Lam NT (2017), PVA had previously been reported as a biocompatible polymer. It was also determined that PVA/CNC_10 was noncytotoxic. In the current study, the cell viability result of PVA/CNC_0, PVA/CNC_10 was above and around 65%, confirming less cytoxicity.\

##### 3.7. PVA_CNC scaffolds enhanced cell proliferation

As PVA_CNC scaffolds showed increased cell viability (Fig. 4), we next determined their effect on cell proliferation. A Muse Cell Analyzer was used to count cell viability in order to observe the effect of bioscaffold on the viability of C _3_H _10_ t _1/2_ cells. Cells were plated in 6-well plates with different scaffold compositions and were incubated for 24 hours at 37C. After 24 hours, cells were trypsinized and viability was checked in a Mouse Cell Analyzer. Fig. 7 (a) shows a representative plot of the percentage of live and dead cell populations excluded from debris in the control and treated groups, and Fig. 7 (b) shows the cell analyser report with different scaffold compositions and control with no scaffold [22]. Cells were analyzed by MUSE cell analyzer after 24 hours of cell plating on bioscaffold. Treatment with PVA/CNC bioscaffold have shown to increase cell viability and enhanced cell growth and also determined that PVA/CNC_10 was non-cytotoxic.

**Fig. 7.**
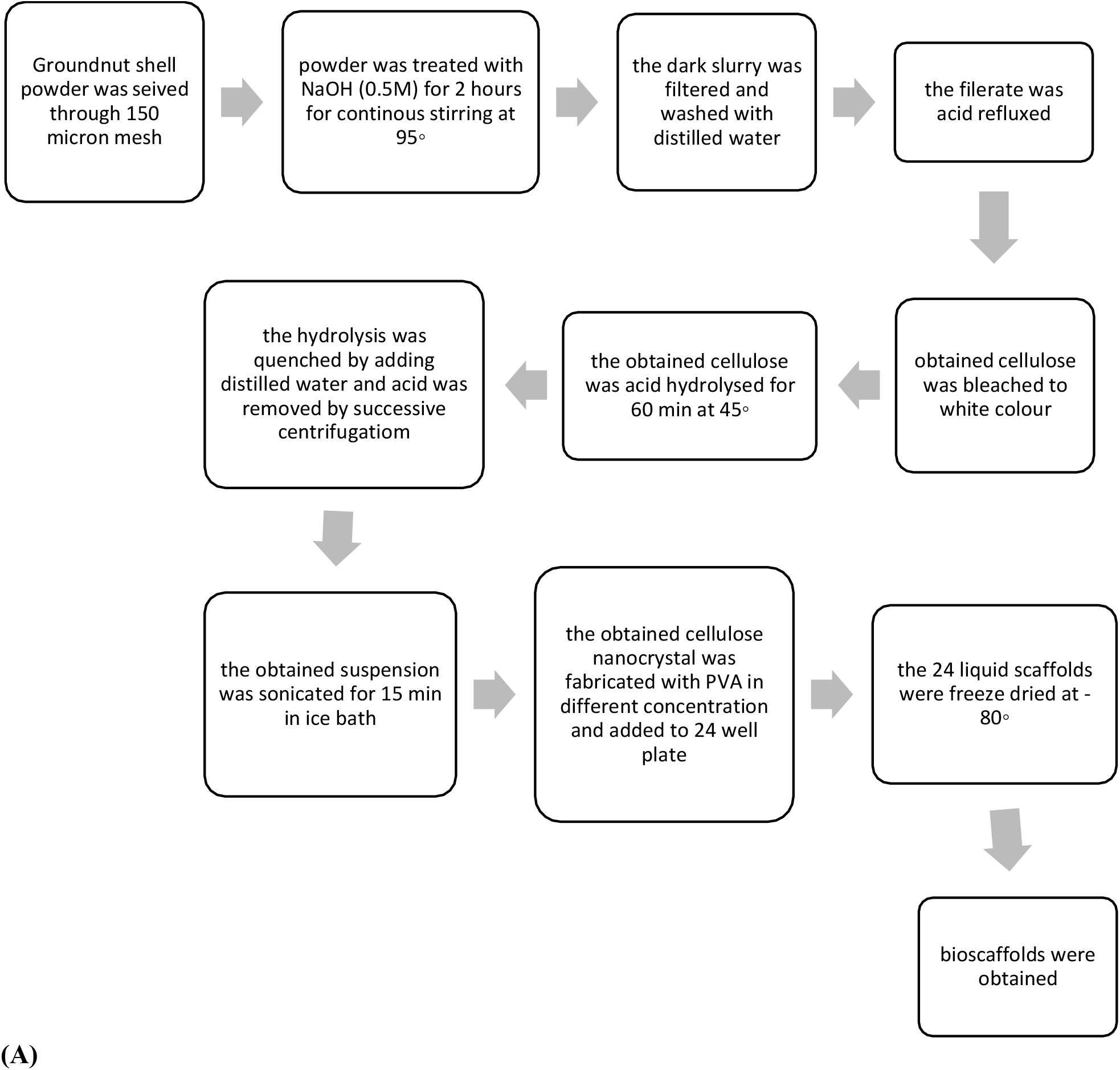
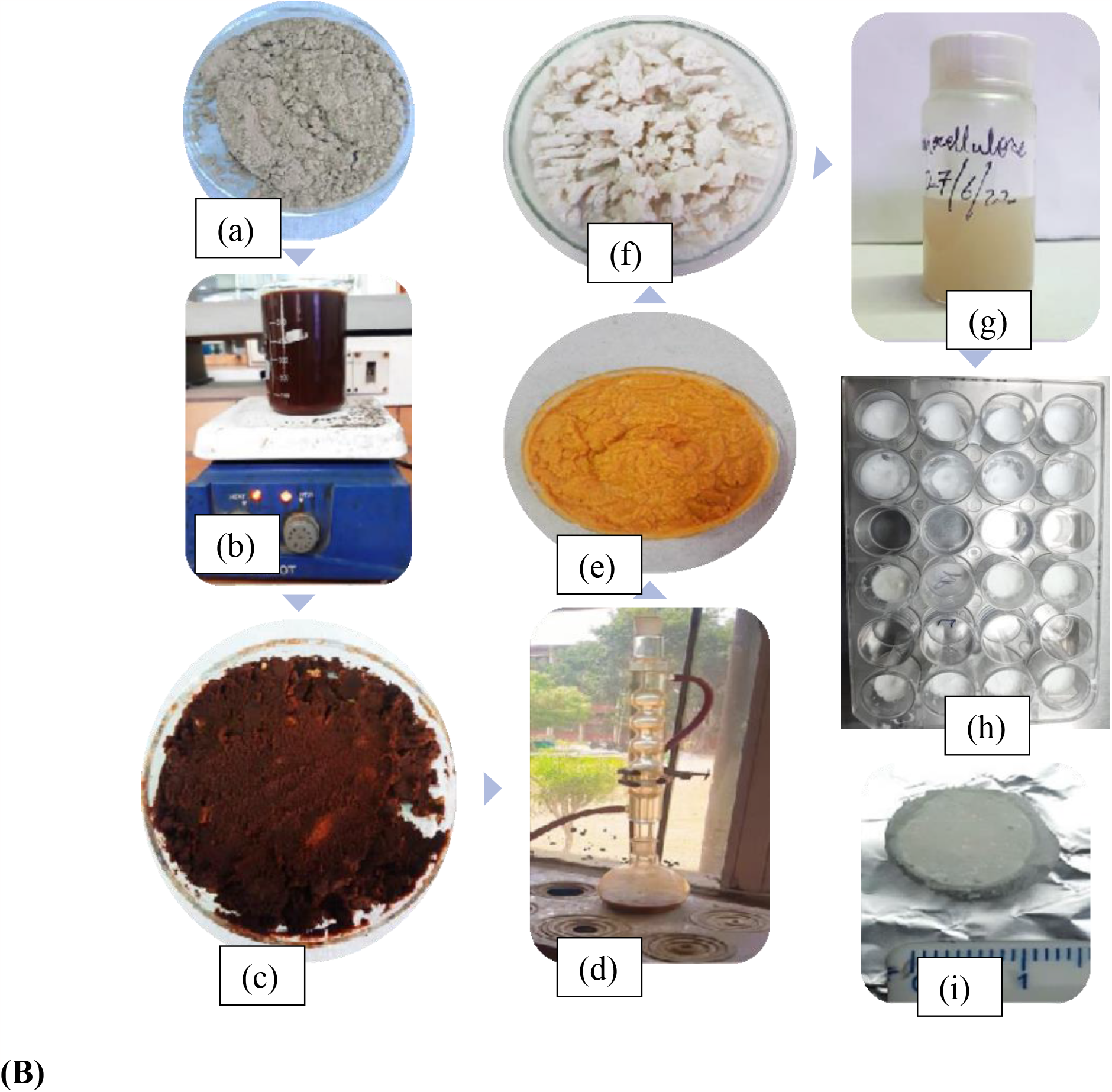
Effect of scaffolds on the proliferation of murine mesenchymal stem cells(C3H10T1/2). After 24 hours of treatment, the cells were then subjected to a cell count and viability assay using a MUSETM automated cell counter. Precise gating adjustments were pre-set to exclude debris, nucleated cells, and live cells from dead cells. (A) representative plot showing the live cell populations and dead cell populations on the left and right quadrants, respectively. Shows the cell viability results of control, PVA/CNC_0, PVA/CNC_5, PVA/CNC_10. PVA was clearly a cell-friendly material with greater than 70% cell viability. (B) Shows the cell analyser report with different scaffold compositions and control with no scaffold [20,23]

##### 3.8. DAPI Staining

The murine mesenchymal stem cell were treated with PVA/CNC_10% for 7 days and were subjected to DAPI staining that showed the presence of adhered and proliferated C_3_H_10_t_½_cells in fig. 8.

**Fig. 8.**
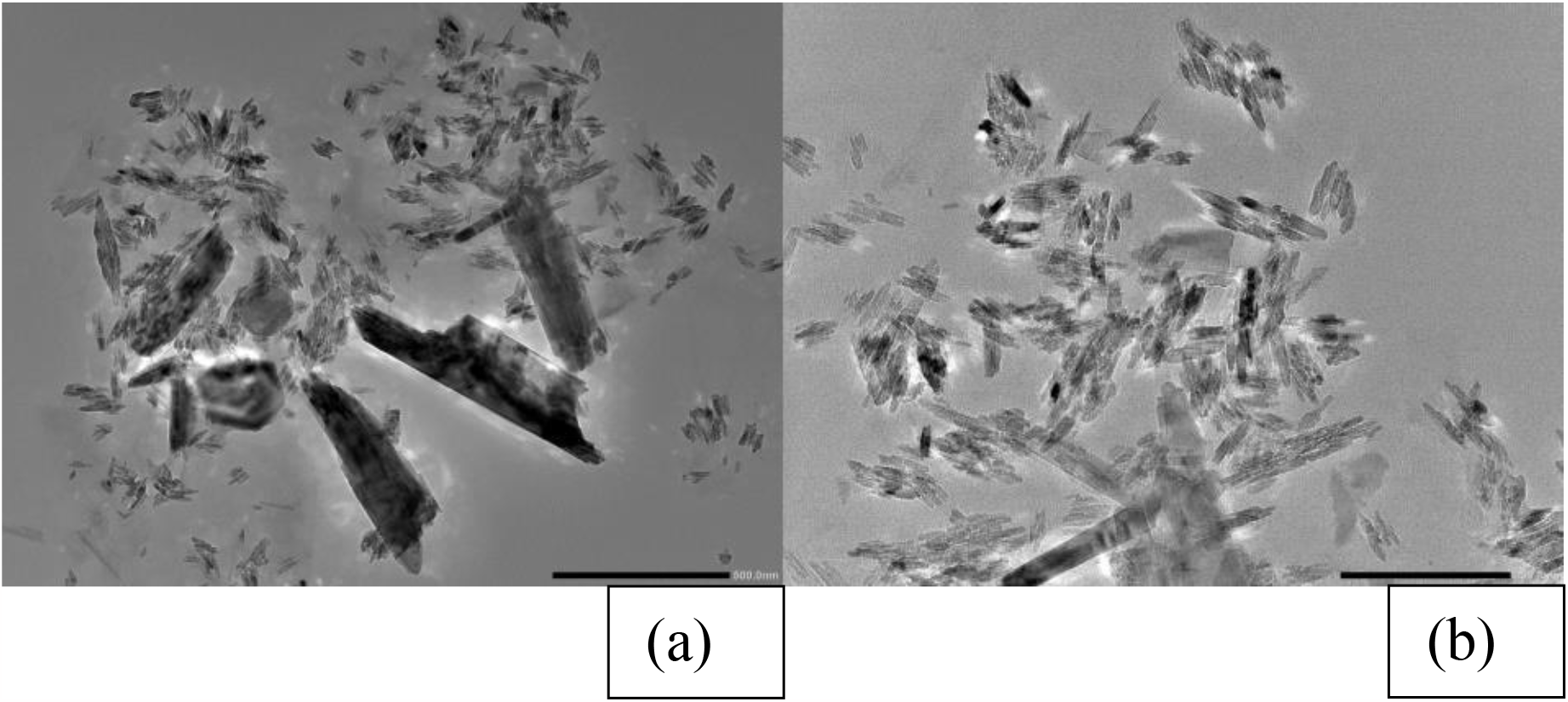
DAPI stain nucleus in scaffold PVA/CNC_10%.

## 4. Conclusion

The present study demonstrated the scaffold activity of peanut shell containing cellulosic components *in vitro*. Peanut shells have a great amount of cellulose fabricated with polyvinyl alcohol to form PVA/CNC composites which used as bioscaffold. The obtained composites were highly porous and the porosity was increased with increasing concentration of CNC in PVA. By treating mesenchymal stem cells with different concentration of fabricated composites viability of mesenchymal stem cells was found to proliferate. Moreover, for C_3_H_10_t_½_ cells, PVA/CNC_10% was less cytototoxic as compared to 0% and 5% concentrations. Our results suggest that treating cells with PVA/CNC_10% the cells got more adhered to composite and proliferated well. A larger study on the effect of growth of various cell lines on the nanocellulose isolated from groundnut shell is needed to confirm these results. In addition, future studies should investigate scaffolding with other components from different waste materials, such as peanut shells. This research helps to elucidate the non-cytotoxic and proliferative effect of peanut shell *in vitro*, further confirming its potential as a functional scaffold for cell tissue engineering.

## 5. Future scope

The scaffold can be utilized for regenerating and restoring damaged tissues and organs.

Scaffold can be further tested for heat tolerance, mechanical compression and swelling behaviour. Future research will have to address PVA/CNC_10% and cell interaction in more detail. Studying refined media compositions containing growth factors and cell diffentiaion factors would also be important to carry out further differentiation of mesenchymal stem cell and its interaction with composite. *In vivo* testing can be done before clinical trials.

## 6. Acknowledgement

The authors are grateful for financial support provided by DBT-Builder Grant vide No. BT/INF/22/SP41295/2020, dated 25/01/2021, sanctioned by Government ofIndia, Ministry of Science and Technology, Department of Biotechnology, New Delhi and Center for Stem Cell Tissue Engineering and Biomedical Excellence, Panjab University, for technical support and facilities provided.

